# Vagally Mediated Heart Rate Variability During REM Sleep is Associated with Retention of Fear Extinction in Generalized Anxiety Disorder

**DOI:** 10.64898/2026.02.02.703359

**Authors:** Cagri Yuksel, Emma McCoy, Lauren Watford, Monami Muranaka, Meltem Sen, Hannah Lax, Lauren Bobowski, Carolina Daffre, Catherine Bostian, Katelyn Oliver, Natasha Lasko, Dan Denis, Edward Pace-Schott

## Abstract

Fear extinction processes are central to the pathology and treatment of anxiety disorders. Emerging evidence implicates rapid eye movement (REM) sleep in the consolidation of extinction memory. Separately, converging theory and empirical work suggest that vagally mediated heart rate variability (VmHRV) serves as a peripheral index of cortico–subcortical regulatory capacity relevant to extinction circuitry. On this basis, we tested the hypothesis that VmHRV during REM sleep would be associated with the retention of extinction memory in individuals with generalized anxiety disorder (GAD). Participants underwent a validated two-day fear conditioning and extinction paradigm. Subjective extinction retention (sERI) was quantified during a recall test 24 hours after learning and ambulatory polysomnography was recorded on the intervening night. As hypothesized, higher REM VmHRV was significantly associated with better extinction retention. This association remained robust after controlling psychotropic medication use and REM density. In contrast, VmHRV during SWS or wakefulness, as well as other REM sleep measures, were not associated with extinction retention. These findings identify REM VmHRV as a significant predictor of extinction memory retention in GAD, extending prior findings in trauma-exposed individuals. We propose that reduced vagal tone indexes compromised prefrontal inhibitory control over amygdala and noradrenergic circuits, thereby impairing REM sleep-dependent consolidation. These results position VmHRV during REM sleep as a potential transdiagnostic biomarker of extinction memory processing and suggest that interventions enhancing vagal tone could improve treatment outcomes in anxiety disorders.

## INTRODUCTION

Fear extinction refers to the process whereby the fear response is gradually reduced after multiple exposures to a conditioned stimulus (CS) without reinforcement. It is central to the contemporary models that explain the core pathology in fear-based anxiety disorders and post-traumatic stress disorder (PTSD) [1-3]. It is also considered to be the mechanism underlying the therapeutic effect of exposure therapy, a modality widely used across psychiatric disorders [4].

Impairments in extinction learning and its retention, along with associated alterations in neural activation patterns, have been consistently reported across various anxiety disorders and PTSD [5-7]. However, the role of extinction processes in generalized anxiety disorder (GAD) remains insufficiently characterized. Unlike anxiety disorders predominantly marked by acute fear responses elicited by specific stimuli or contexts, GAD is defined by generalized worry without an identifiable stimulus or discrete trigger. Consequently, explicit research focused on fear extinction in GAD is sparse, with most available studies grouping participants diagnosed with GAD together with individuals from other anxiety disorders [5], thus limiting the identification of disorder-specific extinction mechanisms.

Despite these limitations, examining extinction processes in GAD is relevant. Current evidence, though limited, suggests that GAD displays impaired extinction and differential engagement of extinction related circuitry [8, 9]. Furthermore, additional indirect evidence from broader anxiety research indicates impaired extinction processes in individuals exhibiting elevated trait anxiety, documented in both adult and pediatric populations [10-12]. Moreover, deficits in fear extinction have increasingly been recognized as a transdiagnostic phenomenon, extending beyond traditionally fear-based disorders to include conditions such as obsessive-compulsive disorder [13, 14] and schizophrenia [15, 16]. Thus, given the use and effectiveness of exposure-based cognitive behavioral therapies in treating GAD [17-19], elucidating extinction processes within this disorder could inform therapeutic interventions and facilitate the identification of predictors for successful treatment outcomes.

Extinction is conceptualized not as the erasure of previously acquired fear responses but rather as the formation of new inhibitory learning, critically dependent upon memory encoding and retrieval processes [1]. Sleep is well established to support memory consolidation [20], with accumulating evidence specifically implicating rapid eye movement sleep (REM) in the consolidation of extinction memories [21, 22, ; however, see 23 indicating a role for non-REM sleep]. In healthy individuals, better extinction recall was associated with the presence of REM during a nap [24], higher overnight %REM [25], less fragmented REM, and increased REM theta power [26]. Additionally, late-night (REM-rich) sleep but not early-night sleep benefited extinction memory [27], and selective REM, but not NREM, deprivation impaired extinction recall [28], suggesting the possibility of a causal role for this sleep stage. Similarly, in trauma-exposed individuals, we recently found that poor extinction recall was associated with lower REM%, shortened REM latency, and increased REM density [22]. In contrast, among individuals with insomnia disorder [26], better extinction recall was associated with less REM%, shorter REM bouts, and longer REM latency. REM abnormalities are widely reported in major depressive disorder and PTSD. The picture in anxiety disorders is less clear, and there are only a few studies that report objective sleep in GAD [29]. Nonetheless, available data indicate REM disturbances, including decreased REM%, shortened or prolonged REM latency, increased REM density, and fewer REM periods [29]. Therefore, it is plausible that REM disruption may be linked with poor memory in GAD.

A specific mechanism that may be associated with the sleep-dependent consolidation of extinction memory is vagal signaling. The vagus is the principal nerve of the parasympathetic nervous system that regulates the activity of most of the internal organs. An influential theory, the neurovisceral integration theory, posits that physiological regulation (i.e., vagal output) and affective regulation (including fear extinction) are closely interrelated self-regulatory processes that rely upon a common cortico-subcortical network [30]. Inhibition by the medial prefrontal cortex within this network allows adaptive and flexible responses to constantly changing environmental demands [30]. Within this framework, vagal output, indexed as vagally mediated heart rate variability (VmHRV), reflects the efficiency of this top-down inhibition. This link is supported by recent empirical evidence, which shows that higher VmHRV during wake is associated with better extinction learning in healthy individuals [31-34].

The vagus is also a predominantly sensory nerve that provides a major bidirectional communication pathway between peripheral organs and the brain. Vagal afferents project to key limbic and cortical regions that are also involved in fear extinction (amygdala, hippocampus, and medial prefrontal cortex [mPFC], among others) [35, 36] and engage ascending multiple neurotransmitter systems, notably norepinephrine, which modulate synaptic plasticity consolidation processes in prefrontal–amygdala–hippocampal circuits [37, 38]. Consistent with this neurobiology, activation of vagal afferents through vagus nerve stimulation induces plasticity in the PFC-amygdala pathway [39] and improves extinction learning and recall in rodents [39-43] and humans [44-46].

Expanding evidence indicates that anxiety disorders, including GAD, are characterized by reduced VmHRV [47]. Given the background summarized above, we hypothesized that reduced VmHRV during REM sleep would be associated with poorer consolidation of extinction memory in GAD, consistent with our recent study, which found a similar link in trauma-exposed individuals [22].

## METHODS

### Participants

This is a secondary analysis of data collected for a study that tested the hypothesis that comorbid insomnia would affect fear extinction in GAD [48]. Thirty-five adults (18–40 years) were recruited from outpatient clinics and community advertisements. Eligibility required meeting DSM-5 criteria for GAD, established via the Structured Clinical Interview for DSM-5, Research Version (SCID-5 RV) administered by an expert psychodiagnostician, and Generalized Anxiety Disorder-7 [GAD-7] score ≥ 10 and Penn State Worry Questionnaire [PSWQ] score ≥ 50. Participants included individuals with none to severe degrees of comorbid DSM-5 insomnia disorder (ID) with symptoms on the Insomnia Severity Index (ISI) ranging from 0 – 25. Other primary sleep disorders were exclusionary. Exclusion criteria also included: current major depressive episode or suicidal ideation; lifetime history of bipolar disorder, psychosis, neurodevelopmental disorder, or neurologic disease likely to affect autonomic or cognitive function; significant medical illness expected to alter sleep or autonomic physiology (e.g., severe cardiovascular disease); recent shift work or trans-meridian travel; current alcohol or drug abuse, positive urine toxicology, or unwillingness to remain alcohol-/drug-free during the protocol; use of specific psychotropic medications such as hypnotics, psychostimulants, antipsychotics, and drugs that influence sweating (e.g., tricyclic antidepressants). All procedures were approved by the Partners Healthcare Institutional Review Board. Participants provided written informed consent and were compensated for participation. Anxiety and sleep symptoms were evaluated using GAD-7, PSWQ, Anxiety Sensitivity Index (ASI), the Pittsburgh Sleep Quality Index (PSQI), and ISI. Of the 35 enrolled participants meeting these criteria, 6 did not complete the study. Demographic and clinical characteristics of the final sample (n=29) are displayed in **Table 1**.

**Table 1.** Demographic and clinical characteristics of the participants (n=29). Data is displayed as n (% of total sample) or mean ± standard deviation (minimum - maximum), where applicable. ASI, Anxiety Sensitivity Index; GAD-7, 7-item Generalized Anxiety Disorder scale; MDD (lifetime), lifetime history of major depressive disorder; OCD, obsessive-compulsive disorder; PSWQ, Penn State Worry Questionnaire; PTSD, posttraumatic stress disorder; SAD, social anxiety disorder; STAI, State-Trait Anxiety Inventory.

|  |  |
| --- | --- |
| **Age** | 24.6±4.2 (19-35) |
| **Sex (%female)** | 27 (93.1%) |
| **Race** |  |
| American Indian<br>or Alaskan Native | 1 (3.4%) |
| Asian | 6 (20.7%) |
| Black or African-American | 2 (6.9%) |
| More than one race | 1 (3.4%) |
| Unknown/unreported | 1 (3.4%) |
| White | 18 (62.1%) |
| **Ethnicity** |  |
| Hispanic or Latino | 4 (13.8%) |
| Not Hispanic or Latino | 24 (82.8%) |
| Unknown/unreported | 1 (3.4%) |
| **Anxiety Symptom Severity** |  |
| ASI | 31.7±9.0 (12-52) |
| GAD-7 | 14.2±3.0 (10-19) |
| PSWQ | 51.9±4.9 (43-63) |
| STAI | 52.2±4.3 (43-60) |
| **Sleep Disturbance Severity** |  |
| ISI | 13.6±6.0 (1-22) |
| PSQI | 8.4±3.6 (1-20) |
| **Medications** |  |
| Antidepressants | 1 (3.4%) |
| Benzodiazepines | 1 (3.4%) |
| **Comorbidities** |  |
| Insomnia | 10 (34.5%) |
| MDD (lifetime) | 12 (41.4%) |
| OCD | 1 (3.4%) |
| Panic Disorder | 2 (6.9%) |
| PTSD | 1 (3.4%) |
| SAD | 6 (20.7%) |
| Simple phobia | 3 (10.3%) |

### Procedures

Following their diagnostic interview or at various times during the 2-week sleep monitoring period described below, participants underwent an acoustic-startle protocol [25, 49-51]. During this protocol, prior to the administration of loud tone startle stimuli, participants completed a 5-minute baseline-monitoring period during which wake electrocardiogram (ECG) was acquired while participants were sitting quietly.

Next, participants completed approximately 2 weeks of home sleep monitoring, during which they continuously wore an actigraph (Actiwatch-2, Philips Respironics) and kept a daily log to document their subjective sleep quality using the Evening-Morning Sleep Questionnaire (EMSQ) [52]. During this period, an acclimation/diagnostic screening night with ambulatory polysomnography (PSG) using the Compumedics Somte-PSG system (Compumedics USA, Inc., Charlotte, NC) was completed to acclimate the participant to PSG and screen for obstructive sleep apnea (OSA) or periodic limb movement disorder (PLMD) symptoms of a severity sufficient to warrant referral for treatment. After this screening/acclimation night, participants completed an additional overnight PSG session that served as the “baseline” sleep recording. Beginning in the evening immediately following the baseline night, participants underwent a two-day fear conditioning and extinction paradigm while undergoing simultaneous fMRI acquisition. Between the two fMRI sessions, participants completed a third overnight ambulatory PSG session, referred to as the “consolidation” night. Full PSG acquisition including ECG and sleep scoring procedures are described in the Supplementary Materials.

### Fear conditioning, extinction learning, and extinction recall

A validated 2-day paradigm [14, 53] was used to probe fear conditioning and extinction during ongoing fMRI recording. This protocol consisted of 4 phases, with Habituation, Fear Conditioning, and Extinction Learning phases taking place on the first day and Extinction Recall 24 hours later. Images of a colored desk lamp (red, yellow, or blue) served as conditioned stimuli (CS). Colored lamps appeared within two different photographic backgrounds (“contexts”) designated as conditioning (office) and extinction (library) contexts. During fear conditioning, a mild electric shock was paired with the presentation of 2 differently colored lamps (CS+) but not a third color (CS-). During extinction learning, one CS+ (CS+E) but not the other (CS+U) was extinguished using presentations without any electric shock.

During each phase, physiologic reactivity for each trial was indexed using skin conductance response (SCR), a measure of sympathetic activity [54]. For all analyses, negative SCRs were coded as zero [55], and then all values were square root transformed. SCRs to the presentation of each of the two CS+s during conditioning (along with their ordinally corresponding CS-), were excluded because pairing with the US, and hence fear learning, had not yet occurred. “Non-conditioners” were defined as those who exhibited less than two non-square-root transformed SCR responses to either of the two CS+s that were equal to or exceeding .05 µS during the Fear Conditioning phase [56]. Six non-conditioners were further excluded from SCR analyses based on these criteria. In addition, SCR data were missing from the Conditioning phase for one participant, from the Recall phase for two participants, and from all phases for one participant, due to recording errors.

Immediately following each phase except Habituation, participants verbally reported shock expectancy for the first and last presentations of each CS (i.e., colored light) appearing in that phase on a scale from 1 (“not expecting a shock at all”) to 5 (“expecting a shock very much”).

Further details of this fear conditioning and extinction protocol are provided in previous publications [26, 57] and in Supplementary Materials.

### Extinction Retention

To examine the association of explicit extinction retention with sleep measures, we used a Subjective Extinction Retention Index (sERI) [22, 26, 58] calculated as: 100 – ((Expectancy to the first CS+E in Extinction Recall/mean expectancy for the last of each CS+ during Fear Conditioning) × 100). Higher sERI values reflect poorer extinction retention (i.e., more persistent expectancy of shock to the extinguished cue). Data on physiological (SCR-based) extinction retention, together with sleep measures, were available for only a small number of participants (n=14) due to non-conditioners and missing SCR data. In addition, our recent work [22] suggested that the physiological REM VmHRV association with extinction retention may be male-specific, whereas the present sample was predominantly female. Accordingly, the main text focuses on associations between sERI and VmHRV. Computation of physiological (SCR-based) extinction retention and its associations with VmHRV and other sleep measures are presented in the Supplementary Materials.

### Heart Rate Variability

HRV was calculated from continuous ECG segments ≥5 minutes in length using Kubios HRV Premium (Kubios Oy, Kuopio, Finland). Prior to analysis, ECG traces were visually inspected for physiological and technical artifacts [59]. Misplaced or ectopic beats were manually corrected if possible. We primarily utilized a “Very Low” (0.45 s) or “Low” (0.35 s) threshold filter (https://www.kubios.com/hrv-preprocessing/) to address any residual outliers. In rare instances where persistent noise exceeded these limits, a “Medium” (0.25 s) filter was applied. Segments were excluded if >7% of total beats required correction/replacement or if noise/non-stationarity precluded reliable estimation. HRV was calculated separately for REM sleep, slow wave sleep (SWS), and wake, as the average of the values across all available segments. High frequency (HF; .15-.4 Hz) absolute power (HF[ms^2^]) was used as an index of vagal activity (VmHRV) because it was shown to be associated with sleep-dependent memory processing [60] and with extinction recall in our recent study [61]. The autoregressive method was used for the analyses, with model order set at 16 [59] and data were transformed by their natural logarithm (lnHF[ms^2^]) for use in all analyses [59]. REM VmHRV during the consolidation night (**Table 2**) was used in the primary analysis that examined the link between VmHRV and extinction retention; VmHRV in other states (SWS and wake) were used in exploratory analyses. Five participants were excluded from the primary analysis, because they did not provide ≥5-minute artifact-free REM segments (n=2) and because of recording errors (n=3).

**Table 2.** Sleep and HRV measures during the consolidation night. Data is displayed as mean ± standard deviation.

|  |  |
| --- | --- |
| **%N1** | 4.3±2.2 |
| **%N2** | 57.1±9.9 |
| **%N3** | 19.4±11.5 |
| **%REM** | 19.3±8.6 |
| **REMD** | 4.8±2.6 |
| **REML** | 105.4±50.0 |
| **REMF** | 13.5±5.8 |
| **REM-HF[ms<sup>2</sup>]** | 879.6±677.2 |
| **SWS-HF[ms<sup>2</sup>]** | 1141.3±1084.3 |

### Sleep Measures

We calculated REM measures that have been shown to be associated with extinction recall [25, 26, 61], including REM%, REM density (REMD), REM latency (REML), REM fragmentation (REMF), and REM theta (REMT), for the consolidation night (**Table 2**). Percent time spent in each sleep stage (%N1-N3, %REM) was computed as a percentage of total sleep time (TST). REMD is the number of rapid eye movements per minute of REM sleep and was calculated from electrooculogram data using an automatic algorithm [62]. REML was calculated as the number of minutes occurring after sleep onset before the first REM epoch. REMF was calculated as the average duration of REM segments [63]. REM segments were defined as continuous REM from the start of at least 1 minute of REM to the onset of at least 1 minute of non-REM or wake [63]. REM theta (REMT) was derived from frontal EEG (F3, F4). For each electrode, we computed power spectral density (PSD) with Welch’s method (5-s windows, 50% overlap) on the first derivative of the EEG to minimize 1/f scaling [64]. Because absolute amplitude varies across individuals due to skull thickness, gyral folding, etc. [64], each spectrum was normalized within participant by dividing power at each frequency by that electrode’s mean total (i.e., 0–30 Hz) power [65, 66]. Theta power was defined as the mean 4–8 Hz power; F3 and F4 were averaged to yield a single REM theta value per participant. Sleep data from three participants were not available due to recording errors.

### Statistical Analyses

#### Verification of fear conditioning and extinction learning

To verify that physiological conditioning and extinction learning occurred, we compared SCRs across the first two (“Early”) and last two (“Late”) trials of each phase (Conditioning and Extinction Learning; [67-76]). As an exception, the first trial in the Conditioning phase was excluded because fear learning had not yet occurred. For each trial, SCR for CS+ was calculated as the average of SCRs for CS+U and to-be CS+E, and SCRs for CS− as the average of their ordinally corresponding CS−s. For both phases, within-subject repeated-measures analyses of variance (rm-ANOVA) were conducted with factors CS type (CS+, CS−), Time (Early, Late), and their interaction. Pre-specified follow-ups were: (i) within-CS time tests (Early to Late) and (ii) planned contrasts of CS+ versus CS− at Late Conditioning, Early Extinction Learning, and Late Extinction Learning.

Similarly, for explicit (expectancy) indices, rm-ANOVAs were conducted with factors CS type (CS+, CS−), Time (first presentation [“First”], last presentation [“Last”]), and their interaction. Expectancy to CS+ during Conditioning was calculated as the average of CS+U and to-be CS+E. Pre-specified follow-ups were: (i) within-CS time tests (First to Last) and (ii) planned contrasts of CS+ versus CS− at the end of Conditioning, and at the onset and end of Extinction Learning.

#### VmHRV reliability and clinical correlates

Night-to-night reliability of REM VmHRV was assessed with intraclass correlation coefficients (ICC; two-way mixed, absolute agreement) between baseline and consolidation nights, and cross-state consistency was examined between REM and wake VmHRV. Associations of VmHRV with demographic and clinical variables were examined to evaluate whether VmHRV was confounded by any of these factors.

#### Association of extinction retention with VmHRV

The primary analysis tested the a priori hypothesis that consolidation-night REM VmHRV would be associated with sERI using bivariate correlation. To evaluate robustness, a follow-up linear regression was conducted with sERI as the dependent variable and REM VmHRV, REM density, and psychotropic medication use as predictors; these covariates were included because they were significantly associated with sERI in a prior study using the same paradigm [22]. Exploratory analyses examined the specificity of any observed association by testing correlations of sERI with VmHRV during SWS and wake, and with other consolidation-night REM measures.

#### General analytic procedures

Shapiro-Wilk tests assessed the normality of each variable’s distribution. Pearson product-moment correlations were used when both variables were normally distributed; Spearman rank-order correlations were used otherwise. For linear regressions, assumptions were evaluated by inspecting residual normality (Q–Q plots), homoscedasticity (residuals vs. fitted values), and influence (Cook’s distance); collinearity was assessed via variance inflation factors (VIF; all values < 2). Outliers exceeding ± 3 standard deviations from the mean were removed. All tests were two-tailed with α = .05. p-values are uncorrected for multiple comparisons unless explicitly noted.

## RESULTS

### Verification of Fear Conditioning and Extinction Learning

#### Skin conductance responses (Figure 1A)

##### Conditioning

The omnibus rm-ANOVA showed significant main effects of CS type (F(1,20) = 11.71, η_*p*_^2^ = 0.37, p = 0.003) and Time (F(1,20) = 25.69, η_*p*_^2^ = 0.56, p < 0.001), with a non-significant CS type × Time interaction (F(1,20) = 1.03, η_*p*_^2^ = 0.05, p = 0.322). Within-CS comparisons indicated decreases from Early to Late subphases for both CS+ (t(20) = 3.96, p < 0.001, d_*z*_ = 0.87; mean difference = 0.27, 95% CI [0.13, 0.41]) and CS− (t(20) = 3.99, p < 0.001, d_*z*_ = 0.87; mean difference = 0.37, 95% CI [0.18, 0.57]). The planned contrast confirmed larger responses to CS+ than CS− at Late Conditioning (t(20) = 3.66, p = 0.002, d_*z*_ = 0.80; mean difference = 0.29, 95% CI [0.12, 0.45]).

**Figure 1.**
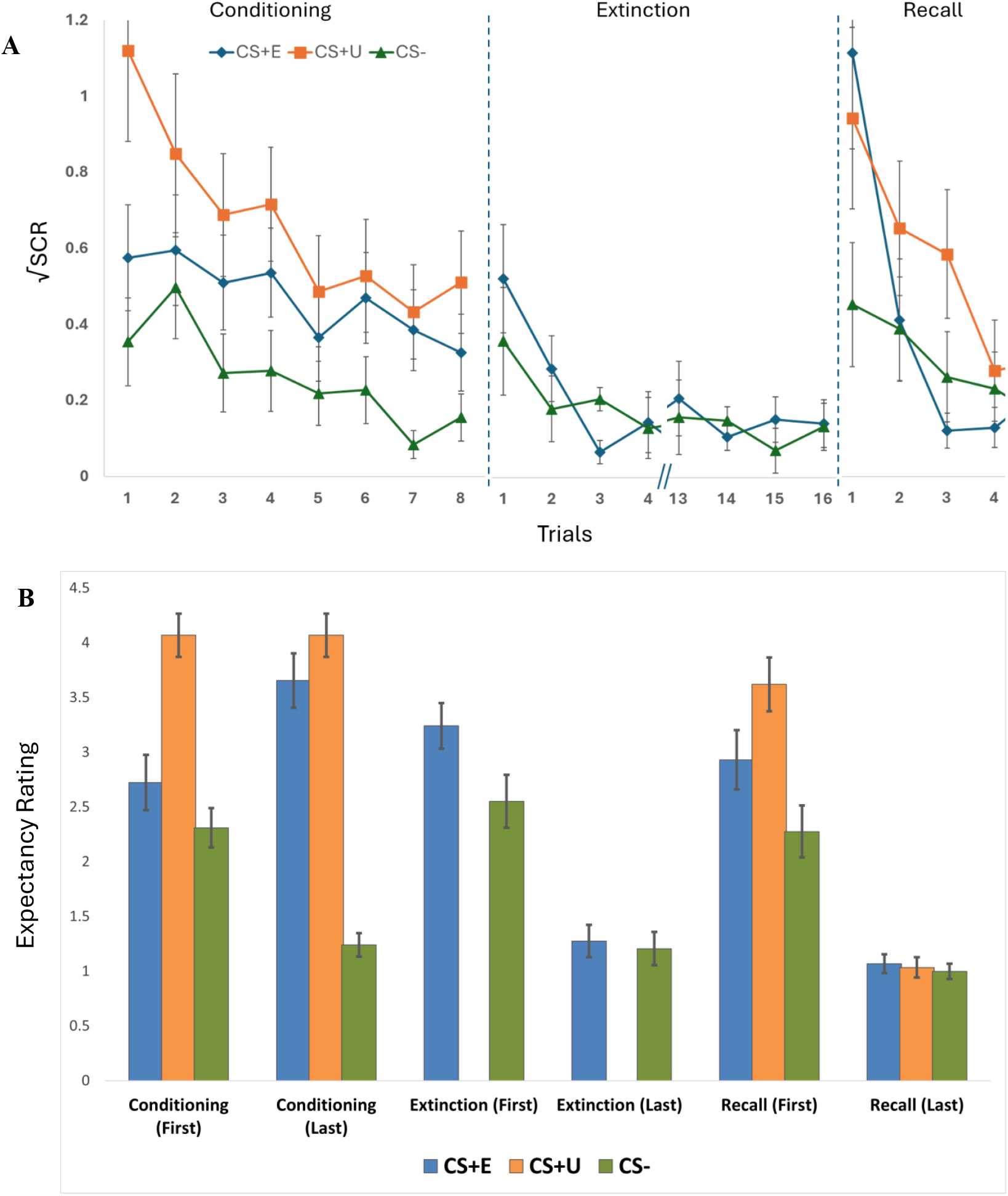
(A) Square-root transformed skin conductance responses (SCR; n = 21) and (B) shock expectancy ratings (n = 29) across Conditioning, Extinction Learning, and Extinction Recall phases. *Conditioning:* Both measures confirmed successful fear acquisition, with significantly greater responses to CS+ (average of CS+E and CS+U) than CS− at late Conditioning. *Extinction Learning:* Both measures demonstrated successful extinction learning. CS+ responses decreased significantly, with CS+ and CS− converging at the end of Extinction. Expectancy ratings showed the expected pattern of CS+ > CS− at Extinction onset before declining to equivalence. SCR did not show significant CS+/CS− differentiation, potentially reflecting lower sensitivity of autonomic measures at this early time point. For display purposes, only the first and last four Extinction trials are shown for SCR; for Recall, only the first four trials are displayed to capture retained extinction memory prior to new extinction learning. Error bars represent standard error.

##### Extinction Learning

The omnibus rm-ANOVA showed a marginal main effect of CS type (F(1,20) = 4.23, η_*p*_^2^ = 0.17, p = 0.053), a significant main effect of Time (F(1,20) = 8.68, η_*p*_^2^ = 0.30, p = 0.008), and no CS type × Time interaction (F(1,20) = 0.64, η_*p*_^2^ = 0.03, p = 0.434). The planned contrast at Early Extinction Learning did not show a significant CS+ versus CS− difference (t(20) = 1.58, p = 0.131, d_*z*_ = 0.34; mean difference = 0.13, 95% CI [−0.04, 0.31]). Within CS+, responses decreased from Early to Late subphases (t(20) = 2.51, p = 0.021, d_*z*_ = 0.55; mean difference = 0.26, 95% CI [0.04, 0.48]). At Late Extinction Learning, CS+ did not differ significantly from CS− (t(20) = 0.92, p = 0.368, d_*z*_ = 0.20; mean difference = 0.05, 95% CI [−0.06, 0.15]).

#### Expectancy ratings (Figure 1B)

##### Conditioning

The omnibus rm-ANOVA showed a significant main effect of CS type (F(1,28) = 214.49, η_*p*_^2^ = 0.88, p < 0.001), a non-significant main effect of Time (F(1,28) = 0.62, η_*p*_^2^ = 0.02, p = 0.437), and a significant CS type × Time interaction (F(1,28) = 36.34, η_*p*_^2^ = 0.56, p < 0.001). Within-CS tests indicated that CS+ ratings increased from the First to the Last presentation (t(28) = −2.76, p = 0.010, d_*z*_ = −0.51; mean difference = −0.78, 95% CI [−1.35, −0.20]), whereas CS− ratings decreased (t(28) = 5.57, p < 0.001, d_*z*_ = 1.04; mean difference = 1.07, 95% CI [0.68, 1.46]). The planned contrast confirmed higher expectancy for CS+ than CS− at the end of Conditioning (t(28) = 13.06, p < 0.001, d_*z*_ = 2.43; mean difference = 2.66, 95% CI [2.24, 3.07]).

##### Extinction Learning

The omnibus rm-ANOVA yielded significant main effects of CS type (F(1,28) = 6.55, η_*p*_^2^ = 0.19, p = 0.016) and Time (F(1,28) = 48.84, η_*p*_^2^ = 0.64, p < 0.001), and a significant CS type × Time interaction (F(1,28) = 6.41, η_*p*_^2^ = 0.19, p = 0.017). CS+ exceeded CS− at the onset of Extinction Learning (t(28) = 2.58, p = 0.016, d_*z*_ = 0.48; mean difference = 0.69, 95% CI [0.14, 1.24]). Within CS+, ratings decreased from the First to the Last presentation (t(28) = 6.95, p < 0.001, d_*z*_ = 1.29; mean difference = 1.97, 95% CI [1.39, 2.54]). At the end of Extinction Learning, CS+ did not differ significantly from CS− (t(28) = 1.44, p = 0.161, d_*z*_ = 0.27; mean difference = 0.07, 95% CI [−0.03, 0.17]).

Collectively, these results demonstrate successful fear conditioning and extinction learning. Expectancy ratings showed the canonical pattern: CS+ exceeded CS− at Extinction Learning onset, declined markedly within CS+ across Extinction Learning, and converged with CS− by Extinction Learning end. For SCR, responses were larger to CS+ than CS− at Late Conditioning, a within-CS+ decline emerged across Extinction Learning, and CS+ and CS− did not differ at Extinction Learning end. SCR did not show a significant CS+ versus CS− difference at Early Extinction Learning, which may reflect lower sensitivity or power for autonomic indices at this early time point.

### VmHRV Reliability and Clinical Correlates

REM VmHRV (lnHF[ms^2^]) showed good night-to-night consistency across the baseline and consolidation nights (ICC [95% CI] = 0.85 [0.58–0.94]). Consolidation night REM VmHRV showed moderate cross-state consistency with wake VmHRV (ICC [95% CI] = 0.75 [0.40– 0.90]). Demographic and clinical variables (age; GAD-7, PSWQ, STAI, ASI, ISI, PSQI) were not significantly correlated with VmHRV during REM on either the consolidation or baseline night, nor with VmHRV during wake.

### Association of Extinction Retention with VmHRV

#### Primary analysis

Consolidation-night REM VmHRV was significantly and inversely correlated with sERI (n = 23, r = −0.53, p = 0.009; **Figure 2**). Because lower sERI reflected better extinction retention, this indicated that higher vagal activity during REM was associated with better extinction retention. When REM density and psychotropic medication use were included as covariates in a follow-up linear regression, the association of REM VmHRV with sERI remained significant (B [SE] = −25.66 [10.49], p = 0.025).

**Figure 2.**
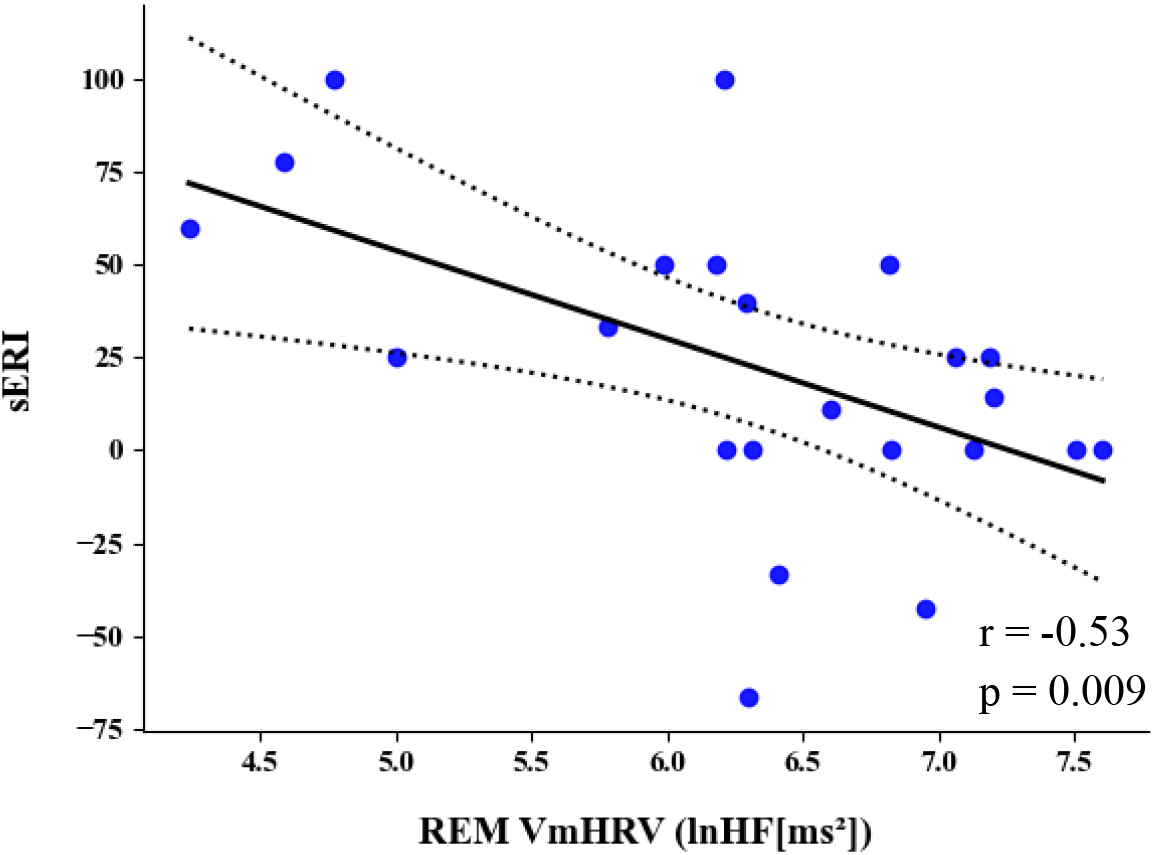
Correlation of sERI with REM VmHRV during the consolidation night. Lower sERI denotes better extinction retention. The dotted lines represent the 95% confidence interval.

#### Exploratory specificity analyses

sERI was not correlated with VmHRV during SWS on the consolidation night (n = 21, r = −0.17, p = 0.450) or wake VmHRV (n = 27, Spearman’s ρ = −0.07, p = 0.726). sERI was also not significantly correlated with other consolidation-night REM variables (REM%, REML, REMD, REMF, REMT), with the exception of a trend-level positive association with REM latency (n = 26, Spearman’s ρ = 0.35, p = 0.079).

## DISCUSSION

In a well-characterized sample of adults with GAD, we found that VmHRV during REM sleep indexed individual differences in subjective extinction retention. Specifically, higher REM VmHRV during the post-extinction learning consolidation night was associated with better retention of extinction memory, as reflected in lower values of the expectancy-based subjective extinction retention index (sERI). This association remained significant when REM density and psychotropic medication use (factors shown to correlate with sERI in a previous study[22]) were included in the model. REM VmHRV showed good night-to-night reliability and was not systematically related to demographic or clinical severity measures, suggesting that it captures a relatively stable physiological characteristic that is not simply a proxy for symptom burden. Exploratory analyses further indicated that sERI was not associated with VmHRV measured during slow-wave sleep or wake, nor with a range of REM-related measures (percentage of REM, REM density, REM fragmentation, REM theta power), aside from a trend-level association with longer REM latency, pointing to a degree of specificity for vagal activity during REM rather than global autonomic tone or conventional REM metrics. Together, these findings implicate REM-related vagal activity as a candidate physiological marker of extinction memory retention in GAD, providing a starting point for integrating autonomic sleep physiology into mechanistic models of anxiety.

The specificity of our results to REM sleep aligns with the body of literature identifying this state as a critical window for the processing of emotionally salient memories in general [77, 78], and, specifically, with the convergent recent evidence implicating its role in the consolidation of fear extinction memory [79, 80]. REM sleep duration and its macro- and micro-structural characteristics have been associated with subsequent extinction recall [22, 25, 26]. Furthermore, selective REM sleep deprivation impairs extinction retention [28], while stimulation of ventromedial prefrontal cortex during REM sleep enhances it [81]. In contrast to the findings in healthy individuals [24-26] as well as clinical samples, including insomnia disorder [26] and trauma-exposed individuals [22], in our GAD sample, we did not find associations with the conventional REM measures. Our findings suggest that, in individuals with clinical anxiety, vagal activity during REM sleep, as indexed by VmHRV, may be a more sensitive indicator of optimal REM sleep that supports the consolidation of fear extinction memory.

Our central finding, that higher VmHRV during REM sleep was associated with better extinction recall, fits within a broader set of studies linking vagal signaling to fear-extinction processes. In healthy individuals, higher VmHRV during wake predicted better safety and extinction learning [31, 32, 34]. Consistent with this, among young individuals with abuse history, VmHRV during wake predicted lower post-traumatic stress disorder symptoms and enhanced extinction learning [33]. In addition to these studies, which show correlations with vagal output, further evidence for vagal involvement in fear extinction comes from studies that engaged vagal afferents. Rodent studies showed that vagal nerve stimulation (VNS) during wake leads to potentiation in the circuitry between infralimbic cortex - basolateral amygdala [39] and consistently improves fear extinction learning [39, 41-43, 82-88], while severing the vagal afferents impairs extinction [89]. In humans also, an expanding body of literature reports that non-invasive VNS enhances extinction learning [44-46, 90-93], including in individuals with specific phobia [92] (however, not all studies find such effects [94-96]). Our results indicate that the link between the vagal pathway and fear extinction extends to the consolidation of extinction memory during sleep. They also dovetail with our recent work [22], which found that higher REM VmHRV predicts better extinction recall above and beyond traditional REM measures, in a large sample of trauma-exposed individuals. Taken together, these findings suggest that REM VmHRV is a cross-diagnostic predictor of extinction retention.

The observed association of VmHRV with extinction retention can be interpreted in light of the neurovisceral integration theory (NVI) [30]. NVI posits that prefrontal cortex mediated top-down inhibition is the common neural substrate for self-regulatory processes, including cognitive, affective and autonomic regulation (e.g., fear extinction and vagal output), and frames VmHRV as a peripheral index of this cortico-subcortical inhibition [30]. From this perspective, the association between lower REM VmHRV and poorer extinction retention likely reflects a functional deficit in mPFC inhibitory control that undermines the consolidation process through two interrelated pathways. First, compromised mPFC function impairs the direct inhibition of the amygdala, the primary neural substrate for the recall of extinction memories [1]. Second, it may be associated with a concurrent failure to downregulate the locus coeruleus–noradrenergic (LC-NA) system, consistent with the recent human neuroimaging and rodent optogenetic studies demonstrating an inverse relationship between VmHRV and LC-NA activity [97, 98]. Such elevated LC-NA has been associated with pathological anxiety and insomnia [99-101], the two defining characteristics of our sample, and is hypothesized to disrupt the specific neurochemical milieu of REM sleep required for the consolidation of emotional memories [79, 102]. Importantly, vagal tone is not immutable. Meta-analytic and experimental evidence demonstrates that behavioral interventions, including heart rate variability biofeedback [103], voluntary slow breathing [104], and physical exercise [105], can enhance VmHRV and provide emotional and cognitive benefits [103, 106-108]. Future research should investigate whether such interventions would improve vagal tone during sleep and improve extinction consolidation.

Several limitations should be considered when interpreting these findings. First, the overall sample was small (n=29) and clinically heterogeneous (e.g., comorbid insomnia and lifetime MDD were common, and a small subset used psychotropic medications), which may limit generalizability and complicate inferences about diagnosis-specific mechanisms. Second, the primary REM VmHRV–sERI association was estimated in a reduced analytic sample (n=23), because participants without sufficient artifact-free REM ECG segments and those with recording errors were necessarily excluded. This missingness could have introduced selection bias, e.g., systematically excluding individuals with more fragmented sleep or noisier physiology. Third, our sample was predominantly female. Although this reflects the higher prevalence of GAD in women, it limits generalizability to males, which is particularly relevant given emerging evidence of sex-specific associations between REM physiology and extinction [22]. Fourth, due to technical constraints and non-conditioners, physiological extinction data (skin conductance) were available for only a subset of participants. Consequently, our primary findings rely on subjective expectancy ratings that may not fully capture implicit autonomic processes. Fifth, the design was correlational and observational. VmHRV was quantified during naturalistic sleep after extinction learning, and therefore the results cannot adjudicate causal direction or isolate whether REM-related VmHRV is tracking extinction-specific consolidation versus broader individual differences in sleep physiology and regulatory capacity.

In conclusion, this study links individual differences in VmHRV during REM sleep to the subsequent retention of extinction memory indexed by an expectancy-based extinction retention measure, in a GAD sample. Taken together with our similar findings in trauma-exposed individuals [22], VmHRV during REM sleep appears to be a transdiagnostic predictor of extinction retention. Interpreted through neurovisceral integration theory, the association is consistent with the possibility that lower REM VmHRV indexes reduced prefrontal top-down inhibitory control, with downstream consequences for amygdala and LC–NA regulation that impairs extinction-memory consolidation. While the design does not support causal claims, the results add to converging evidence that autonomic physiology and sleep-state context are relevant to extinction-related processes and motivate future studies that evaluate whether interventions that target vagal tone during sleep are accompanied by improvements in extinction retention in clinical samples.

## References

1. Milad, M.R. and G.J. Quirk, Fear extinction as a model for translational neuroscience: ten years of progress. Annu Rev Psychol, 2012. 63: p. 129–51.

2. Zuj, D.V., et al., The centrality of fear extinction in linking risk factors to PTSD: A narrative review. Neurosci Biobehav Rev, 2016. 69: p. 15–35.

3. Milad, M.R., B.L. Rosenbaum, and N.M. Simon, Neuroscience of fear extinction: implications for assessment and treatment of fear-based and anxiety related disorders. Behav Res Ther, 2014. 62: p. 17–23.

4. Craske, M.G., et al., Maximizing exposure therapy: an inhibitory learning approach. Behav Res Ther, 2014. 58: p. 10–23.

5. Marin, M.F., et al., Skin Conductance Responses and Neural Activations During Fear Conditioning and Extinction Recall Across Anxiety Disorders. JAMA Psychiatry, 2017. 74(6): p. 622–631.

6. Duits, P., et al., Updated meta-analysis of classical fear conditioning in the anxiety disorders. Depress Anxiety, 2015. 32(4): p. 239–53.

7. Graham, B.M. and M.R. Milad, The study of fear extinction: implications for anxiety disorders. Am J Psychiatry, 2011. 168(12): p. 1255–65.

8. Pitman, R.K. and S.P. Orr, Test of the conditioning model of neurosis: differential aversive conditioning of angry and neutral facial expressions in anxiety disorder patients. J Abnorm Psychol, 1986. 95(3): p. 208–13.

9. Marin, M.F., et al., Multimodal Categorical and Dimensional Approaches to Understanding Threat Conditioning and Its Extinction in Individuals With Anxiety Disorders. JAMA Psychiatry, 2020. 77(6): p. 618–627.

10. Gazendam, F.J., J.H. Kamphuis, and M. Kindt, Deficient safety learning characterizes high trait anxious individuals. Biol Psychol, 2013. 92(2): p. 342–52.

11. Liberman, L.C., et al., Evidence for retarded extinction of aversive learning in anxious children. Behav Res Ther, 2006. 44(10): p. 1491–502.

12. Dibbets, P., A. van den Broek, and E.A. Evers, Fear conditioning and extinction in anxiety- and depression-prone persons. Memory, 2015. 23(3): p. 350–64.

13. McLaughlin, N.C., et al., Extinction retention and fear renewal in a lifetime obsessive-compulsive disorder sample. Behav Brain Res, 2015. 280: p. 72–7.

14. Milad, M.R., et al., Deficits in conditioned fear extinction in obsessive-compulsive disorder and neurobiological changes in the fear circuit. JAMA Psychiatry, 2013. 70(6): p. 608–18; quiz 554.

15. Holt, D.J., et al., Extinction memory is impaired in schizophrenia. Biol Psychiatry, 2009. 65(6): p. 455–63.

16. Holt, D.J., et al., Failure of neural responses to safety cues in schizophrenia. Arch Gen Psychiatry, 2012. 69(9): p. 893–903.

17. Sars, D. and A. van Minnen, On the use of exposure therapy in the treatment of anxiety disorders: a survey among cognitive behavioural therapists in the Netherlands. BMC Psychol, 2015. 3(1): p. 26.

18. Hoyer, J., et al., Worry exposure versus applied relaxation in the treatment of generalized anxiety disorder. Psychother Psychosom, 2009. 78(2): p. 106–15.

19. Berg, H., et al., A randomized clinical trial of behavioral activation and exposure-based therapy for adults with generalized anxiety disorder. J Mood Anxiety Disord, 2023. 1.

20. Klinzing, J.G., N. Niethard, and J. Born, Mechanisms of systems memory consolidation during sleep. Nat Neurosci, 2019. 22(10): p. 1598–1610.

21. Davidson, P. and E. Pace-Schott, The role of sleep in fear learning and memory. Curr Opin Psychol, 2020. 34: p. 32–36.

22. Yuksel, C., et al., REM disruption and REM vagal activity predict extinction recall in trauma-exposed individuals. Psychol Med, 2024: p. 1–12.

23. Denis, D., et al., Slow oscillation-sleep spindle coupling is associated with expectancy measures of fear extinction retention in trauma-exposed individuals. Biol Psychiatry Cogn Neurosci Neuroimaging, 2025.

24. Spoormaker, V.I., et al., The neural correlates and temporal sequence of the relationship between shock exposure, disturbed sleep and impaired consolidation of fear extinction. J Psychiatr Res, 2010. 44(16): p. 1121–8.

25. Pace-Schott, E.F., et al., Interactions of time of day and sleep with between-session habituation and extinction memory in young adult males. Exp Brain Res, 2014. 232(5): p. 1443–58.

26. Bottary, R., et al., Fear extinction memory is negatively associated with REM sleep in insomnia disorder. Sleep, 2020. 43(7).

27. Menz, M.M., J.S. Rihm, and C. Buchel, REM Sleep Is Causal to Successful Consolidation of Dangerous and Safety Stimuli and Reduces Return of Fear after Extinction. J Neurosci, 2016. 36(7): p. 2148–60.

28. Spoormaker, V.I., et al., Effects of rapid eye movement sleep deprivation on fear extinction recall and prediction error signaling. Hum Brain Mapp, 2012. 33(10): p. 2362–76.

29. Cox, R.C. and B.O. Olatunji, Sleep in the anxiety-related disorders: A meta-analysis of subjective and objective research. Sleep Med Rev, 2020. 51: p. 101282.

30. Thayer, J.F., et al., Heart rate variability, prefrontal neural function, and cognitive performance: the neurovisceral integration perspective on self-regulation, adaptation, and health. Ann Behav Med, 2009. 37(2): p. 141–53.

31. Pappens, M., et al., Resting heart rate variability predicts safety learning and fear extinction in an interoceptive fear conditioning paradigm. PLoS One, 2014. 9(9): p. e105054.

32. Wendt, J., et al., Resting heart rate variability is associated with inhibition of conditioned fear. Psychophysiology, 2015. 52(9): p. 1161–6.

33. Jenness, J.L., et al., Extinction Learning as a Potential Mechanism Linking High Vagal Tone with Lower PTSD Symptoms among Abused Youth. J Abnorm Child Psychol, 2019. 47(4): p. 659–670.

34. Wendt, J., et al., Vagally mediated heart rate variability and safety learning: Effects of instructions and number of extinction trials. Psychophysiology, 2019. 56(10): p. e13404.

35. Neuhuber, W.L. and H.R. Berthoud, Functional anatomy of the vagus system - Emphasis on the somato-visceral interface. Auton Neurosci, 2021. 236: p. 102887.

36. Suarez, A.N., et al., Gut vagal sensory signaling regulates hippocampus function through multi-order pathways. Nat Commun, 2018. 9(1): p. 2181.

37. Roosevelt, R.W., et al., Increased extracellular concentrations of norepinephrine in cortex and hippocampus following vagus nerve stimulation in the rat. Brain Res, 2006. 1119(1): p. 124–32.

38. McGaugh, J.L., Making lasting memories: remembering the significant. Proc Natl Acad Sci U S A, 2013. 110 Suppl 2(Suppl 2): p. 10402–7.

39. Pena, D.F., et al., Vagus nerve stimulation enhances extinction of conditioned fear and modulates plasticity in the pathway from the ventromedial prefrontal cortex to the amygdala. Front Behav Neurosci, 2014. 8: p. 327.

40. Pena, D.F., N.D. Engineer, and C.K. McIntyre, Rapid remission of conditioned fear expression with extinction training paired with vagus nerve stimulation. Biol Psychiatry, 2013. 73(11): p. 1071–7.

41. Noble, L.J., et al., Effects of vagus nerve stimulation on extinction of conditioned fear and post-traumatic stress disorder symptoms in rats. Transl Psychiatry, 2017. 7(8): p. e1217.

42. Noble, L.J., et al., Vagus nerve stimulation promotes generalization of conditioned fear extinction and reduces anxiety in rats. Brain Stimul, 2019. 12(1): p. 9–18.

43. Souza, R.R., et al., Timing of vagus nerve stimulation during fear extinction determines efficacy in a rat model of PTSD. Sci Rep, 2022. 12(1): p. 16526.

44. Szeska, C., et al., Promoting long-term inhibition of human fear responses by noninvasive transcutaneous vagus nerve stimulation during extinction training. Sci Rep, 2020. 10(1): p. 1529.

45. Burger, A.M., et al., The effect of transcutaneous vagus nerve stimulation on fear generalization and subsequent fear extinction. Neurobiol Learn Mem, 2019. 161: p. 192–201.

46. Burger, A.M., et al., The effects of transcutaneous vagus nerve stimulation on conditioned fear extinction in humans. Neurobiol Learn Mem, 2016. 132: p. 49–56.

47. Wang, Z., et al., Heart rate variability in generalized anxiety disorder, major depressive disorder and panic disorder: A network meta-analysis and systematic review. J Affect Disord, 2023. 330: p. 259–266.

48. Seo, J., et al., Local and network neural activations and their associations with sleep parameters during threat conditioning and extinction in persons with Generalized Anxiety Disorder with and without Insomnia Disorder. bioRxiv, 2025.

49. Carson, M.A., et al., Physiologic reactivity to startling tones in female Vietnam nurse veterans with PTSD. Journal of Traumatic Stress, 2007. 20(5): p. 657–66.

50. Buhlmann, U., et al., Physiologic responses to loud tones in individuals with obsessive-compulsive disorder. Psychosomatic Medicine, 2007. 69(2): p. 166–72.

51. Mueller-Pfeiffer, C., et al., Cortical and cerebellar modulation of autonomic responses to loud sounds. Psychophysiology, 2014. 51(1): p. 60–9.

52. Pace-Schott, E.F., et al., Nightcap measurement of sleep quality in self-described good and poor sleepers. Sleep, 1994. 17(8): p. 688–92.

53. Milad, M.R., et al., Recall of fear extinction in humans activates the ventromedial prefrontal cortex and hippocampus in concert. Biol Psychiatry, 2007. 62(5): p. 446–54.

54. Dawson, M.E., A.M. Schell, and D.L. Filion, The electrodermal system. In: Cacioppo JT, Tassinary LG, Berntson GG, eds. Handbook of Psychophysiology. Third ed. 2007, New York, NY: Cambridge University Press.

55. Lonsdorf, T.B., et al., Don’t fear ‘fear conditioning’: Methodological considerations for the design and analysis of studies on human fear acquisition, extinction, and return of fear. Neurosci Biobehav Rev, 2017. 77: p. 247–285.

56. Orr, S.P., et al., De novo conditioning in trauma-exposed individuals with and without posttraumatic stress disorder. J Abnorm Psychol, 2000. 109(2): p. 290–8.

57. Pace-Schott, E.F., et al., Sleep promotes generalization of extinction of conditioned fear. Sleep, 2009. 32(1): p. 19–26.

58. Yuksel, C., et al., REM disruption and REM vagal activity predict extinction recall in trauma-exposed individuals. Psychol Med, 2024. 54(16): p. 1–12.

59. Laborde, S., E. Mosley, and J.F. Thayer, Heart Rate Variability and Cardiac Vagal Tone in Psychophysiological Research - Recommendations for Experiment Planning, Data Analysis, and Data Reporting. Front Psychol, 2017. 8: p. 213.

60. Whitehurst, L.N., et al., Autonomic activity during sleep predicts memory consolidation in humans. Proc Natl Acad Sci U S A, 2016. 113(26): p. 7272–7.

61. Yuksel, C., et al., REM disruption and REM Vagal Activity Predict Extinction Recall in Trauma-Exposed Individuals. bioRxiv, 2024.

62. Yetton, B.D., et al., Automatic detection of rapid eye movements (REMs): A machine learning approach. J Neurosci Methods, 2016. 259: p. 72–82.

63. Mellman, T.A., et al., REM sleep and the early development of posttraumatic stress disorder. Am J Psychiatry, 2002. 159(10): p. 1696–701.

64. Cox, R., et al., Individual Differences in Frequency and Topography of Slow and Fast Sleep Spindles. Front Hum Neurosci, 2017. 11: p. 433.

65. Denis, D., et al., Beta spectral power during sleep is associated with impaired recall of extinguished fear. Sleep, 2023. 46(10).

66. Cunningham, T.J., et al., Sleep spectral power correlates of prospective memory maintenance. Learn Mem, 2021. 28(9): p. 291–299.

67. Craske, M.G., et al., Is aversive learning a marker of risk for anxiety disorders in children? Behav Res Ther, 2008. 46(8): p. 954–67.

68. Gerlicher, A.M.V., V.N. Metselaar, and M. Kindt, In search of the behavioral effects of fear: A paradigm to assess conditioned suppression in humans. Psychophysiology, 2022. 59(10): p. e14079.

69. Merz, C.J., A. Eichholtz, and O.T. Wolf, Acute stress reduces out-group related safety signaling during fear reinstatement in women. Sci Rep, 2020. 10(1): p. 2092.

70. Hornstein, E.A., et al., Social support and fear-inhibition: an examination of underlying neural mechanisms. Soc Cogn Affect Neurosci, 2024. 19(1).

71. Meir Drexler, S. and O.T. Wolf, Stress disrupts the reconsolidation of fear memories in men. Psychoneuroendocrinology, 2017. 77: p. 95–104.

72. Meir Drexler, S., et al., Cortisol effects on fear memory reconsolidation in women. Psychopharmacology (Berl), 2016. 233(14): p. 2687–97.

73. Martinez, K.G., et al., Correlations between psychological tests and physiological responses during fear conditioning and renewal. Biol Mood Anxiety Disord, 2012. 2: p. 16.

74. Pattwell, S.S., et al., Altered fear learning across development in both mouse and human. Proc Natl Acad Sci U S A, 2012. 109(40): p. 16318–23.

75. Coelho, C.A., J.E. Dunsmoor, and E.A. Phelps, Compound stimulus extinction reduces spontaneous recovery in humans. Learn Mem, 2015. 22(12): p. 589–93.

76. Golkar, A., et al., Are fear memories erasable?-reconsolidation of learned fear with fear-relevant and fear-irrelevant stimuli. Front Behav Neurosci, 2012. 6: p. 80.

77. Schafer, S.K., et al., Sleep’s impact on emotional recognition memory: A meta-analysis of whole-night, nap, and REM sleep effects. Sleep Med Rev, 2020. 51: p. 101280.

78. Yuksel, C., et al., Both slow wave and rapid eye movement sleep contribute to emotional memory consolidation. Commun Biol, 2025. 8(1): p. 485.

79. Pace-Schott, E.F., J. Seo, and R. Bottary, The influence of sleep on fear extinction in trauma-related disorders. Neurobiol Stress, 2023. 22: p. 100500.

80. Bottary, R., L.D. Straus, and E.F. Pace-Schott, The Impact of Sleep on Fear Extinction. Curr Top Behav Neurosci, 2023. 64: p. 133–156.

81. Marković, V., et al., Plasticity induction in the ventromedial prefrontal cortex during REM sleep improves fear extinction memory consolidation. bioRxiv, 2025: p. 2025.03.31.646182.

82. Noble, L.J., et al., Peripheral effects of vagus nerve stimulation on anxiety and extinction of conditioned fear in rats. Learn Mem, 2019. 26(7): p. 245–251.

83. Souza, R.R., et al., Vagus nerve stimulation reverses the extinction impairments in a model of PTSD with prolonged and repeated trauma. Stress, 2019. 22(4): p. 509–520.

84. Souza, R.R., et al., Efficient parameters of vagus nerve stimulation to enhance extinction learning in an extinction-resistant rat model of PTSD. Prog Neuropsychopharmacol Biol Psychiatry, 2020. 99: p. 109848.

85. Souza, R.R., et al., Vagus nerve stimulation enhances fear extinction as an inverted-U function of stimulation intensity. Exp Neurol, 2021. 341: p. 113718.

86. Alvarez-Dieppa, A.C., et al., Vagus Nerve Stimulation Enhances Extinction of Conditioned Fear in Rats and Modulates Arc Protein, CaMKII, and GluN2B-Containing NMDA Receptors in the Basolateral Amygdala. Neural Plast, 2016. 2016: p. 4273280.

87. Alvarez-Dieppa, A.C., et al., Vagus nerve stimulation rescues impaired fear extinction and social interaction in a rat model of autism spectrum disorder. J Affect Disord, 2025. 374: p. 505–512.

88. Calderon-Williams, D.R., et al., Optogenetic inhibition of the locus coeruleus blocks vagus nerve stimulation-induced enhancement of extinction of conditioned fear in rats. Learn Mem, 2024. 31(12).

89. Klarer, M., et al., Gut vagal afferents differentially modulate innate anxiety and learned fear. J Neurosci, 2014. 34(21): p. 7067–76.

90. Burger, A.M., et al., Mixed evidence for the potential of non-invasive transcutaneous vagal nerve stimulation to improve the extinction and retention of fear. Behav Res Ther, 2017. 97: p. 64–74.

91. Szeska, C., et al., Attentive immobility in the face of inevitable distal threat-Startle potentiation and fear bradycardia as an index of emotion and attention. Psychophysiology, 2021. 58(6): p. e13812.

92. Szeska, C., et al., Ready for translation: non-invasive auricular vagus nerve stimulation inhibits psychophysiological indices of stimulus-specific fear and facilitates responding to repeated exposure in phobic individuals. Transl Psychiatry, 2025. 15(1): p. 135.

93. Zhang, X., et al., Transcutaneous vagus nerve stimulation modulates fear memory extinction and neural responses in humans. Cereb Cortex, 2025. 35(8).

94. Burger, A.M., et al., Transcutaneous vagus nerve stimulation and extinction of prepared fear: A conceptual non-replication. Sci Rep, 2018. 8(1): p. 11471.

95. D’Agostini, M., et al., Transcutaneous Auricular Vagus Nerve Stimulation Does Not Accelerate Fear Extinction: A Randomized, Sham-Controlled Study. Psychophysiology, 2025. 62(1): p. e14754.

96. Genheimer, H., et al., Reinstatement of contextual conditioned anxiety in virtual reality and the effects of transcutaneous vagus nerve stimulation in humans. Sci Rep, 2017. 7(1): p. 17886.

97. Mather, M., et al., Higher locus coeruleus MRI contrast is associated with lower parasympathetic influence over heart rate variability. Neuroimage, 2017. 150: p. 329–335.

98. Wang, X., et al., Optogenetic stimulation of locus ceruleus neurons augments inhibitory transmission to parasympathetic cardiac vagal neurons via activation of brainstem alpha1 and beta1 receptors. J Neurosci, 2014. 34(18): p. 6182–9.

99. Van Someren, E.J.W., Brain mechanisms of insomnia: new perspectives on causes and consequences. Physiol Rev, 2021. 101(3): p. 995–1046.

100. Morris, L.S., et al., The role of the locus coeruleus in the generation of pathological anxiety. Brain Neurosci Adv, 2020. 4: p. 2398212820930321.

101. Daffre, C., et al., Rapid eye movement sleep parasympathetic activity predicts wake hyperarousal symptoms following a traumatic event. J Sleep Res, 2023. 32(1): p. e13685.

102. Cabrera, Y., et al., Overnight neuronal plasticity and adaptation to emotional distress. Nat Rev Neurosci, 2024. 25(4): p. 253–271.

103. Lehrer, P., et al., Heart Rate Variability Biofeedback Improves Emotional and Physical Health and Performance: A Systematic Review and Meta Analysis. Appl Psychophysiol Biofeedback, 2020. 45(3): p. 109–129.

104. Laborde, S., et al., Effects of voluntary slow breathing on heart rate and heart rate variability: A systematic review and a meta-analysis. Neurosci Biobehav Rev, 2022. 138: p. 104711.

105. El-Malahi, O., et al., Beneficial impacts of physical activity on heart rate variability: A systematic review and meta-analysis. PLoS One, 2024. 19(4): p. e0299793.

106. Tinello, D., M. Kliegel, and S. Zuber, Does Heart Rate Variability Biofeedback Enhance Executive Functions Across the Lifespan? A Systematic Review. J Cogn Enhanc, 2022. 6(1): p. 126–142.

107. Lee, S.H., D.S. Park, and C.H. Song, The Effect of Deep and Slow Breathing on Retention and Cognitive Function in the Elderly Population. Healthcare (Basel), 2023. 11(6).

108. Schwarck, S., et al., Heart Rate Variability During Physical Exercise Is Associated With Improved Cognitive Performance in Alzheimer’s Dementia Patients-A Longitudinal Feasibility Study. Front Sports Act Living, 2021. 3: p. 684089.

