## Supplemental Materials for "Vagally Mediated Heart Rate Variability During REM Sleep is Associated with Retention of Fear Extinction in Generalized Anxiety Disorder"

**Fear Conditioning, Extinction Learning, and Extinction Recall**

A validated 2-day paradigm [1, 2] was used to probe fear conditioning and extinction during ongoing fMRI recording. This protocol consisted of 4 phases, with Habituation, Fear Conditioning, and Extinction Learning phases taking place on the first day and Extinction Recall 24 hours later. During each phase, images of a colored desk lamp (red, yellow, or blue) appearing in a contextual background (office for conditioning context and conference room for extinction context) served as conditioned stimuli (CS). Context images were presented for nine seconds, with three seconds with the lamp off and six seconds with the lamp on (red, yellow or blue). The unconditioned stimulus (US) was a mild (0.8-4.0 mA), 500 msec electric shock delivered to the index and middle fingers of participants' right hand using a Coulbourn Transcutaneous Aversive Finger Stimulator (Coulbourn Instruments, Allentown, PA). Prior to entering the scanner, participants were administered increasing intensities of shock and they each selected a level that they perceived as "highly annoying but not painful". Prior to entering the scanner, MRI-safe 11-mm, Ag/AgCl skin conductance monitoring electrodes were attached to the participant's left palm. Skin conductance level (SCL) was continuously monitored at 37.5 Hz using the MP150 system with Acqknowledge 4.3 (BIOPAC Systems, Inc., Goleta, CA) software.

During Habituation, all six possible combinations of lamp colors and contexts were presented across six trials. During the following Fear Conditioning phase, two of the three colored lamps (CS+) were each presented 8 times paired with the US at stimulus offset, on a partial reinforcement schedule (5 out of 8 presentations were paired with US). The third lamp color, which was never paired with US (CS-), was interspersed among the CS+s for a total of 16 presentations. Fear Conditioning was followed by Extinction Learning, during which one CS+ (CS+E) was presented in the extinction context 16 times without the US along with 16 interspersed presentations of the CS-. The other CS+ remained conditioned but unextinguished (CS+U). During Extinction Recall, which took place 24 hours later, each CS+ was presented 8 times in the extinction context, with no US, along with 16 interspersed CS-.

During each phase, physiologic reactivity for each trial was indexed using skin conductance response (SCR), a measure of sympathetic activity [3], calculated as the mean SCL in microSiemens (μS) during the last 2 seconds of context presentation subtracted from the maximum SCL during the 6 seconds of CS presentation. Negative SCRs were coded as zero [4], and then all values were square root transformed. "Non-conditioners" were defined as those who exhibited less than two non-square-root transformed SCR responses to either of the two CS+s that were equal to or exceeding .05 μS during the Fear Conditioning phase [5].

Immediately following each phase except Habituation, participants verbally reported shock expectancy for the first and last presentations of each CS (i.e., colored light) appearing in that phase on a scale from 1 ("not expecting a shock at all") to 5 ("expecting a shock very much").

Ratings were obtained at this frequency with the goal of capturing the shift in expectancy from baseline to the end of the corresponding session.

### Ambulatory Polysomnography

Ambulatory polysomnography (PSG) was recorded on 3 nights using the Somte-PSG ambulatory sleep monitor (Compumedics USA, Charlotte, NC, USA). Sampling rate was 256 Hz. EEG data were acquired using six EEG channels (F3, F4, C3, C4, O1, O2; positioned according to the 10-20 system). Additional electrodes were placed on bilateral mastoids, above the right and below the left eye (EOG), under the chin (EMG), and below the right clavicle and in the left fifth intercostal space (ECG). Participants returned home to sleep after being instrumented. During the acclimation/screening (first) PSG night, additional channels for pulse-oximeter, respiration transducer belts, nasal cannula, and tibialis movement sensors were added to screen for obstructive sleep apnea (OSA) and Periodic Limb Movement Disorder (PLMD). All sleep records were scored by an experienced, registered polysomnographic technologist according to American Academy of Sleep Medicine criteria [6].

### Physiological Extinction Retention

A physiological extinction retention index (ERI) was calculated using the following formula: (Average of the SCRs of the first 4 CS+E presentations at Extinction Recall phase/maximum SCR to the "to-be" CS+E during the Fear Conditioning phase) x 100) [7]. Only the first 4 CS+E trials from the Extinction Recall phase were included in this calculation in order to avoid confounding recalled extinction with new extinction learning. Higher ERI reflected lower extinction memory.

There were no significant correlations between ERI and rapid eye movement sleep (REM) vagally-mediated heart rate variability (VmHRV; lnHF[$ms^2$]) during the consolidation night ($n=13$, Spearman's $\rho = 0.06$, $p = 0.845$). ERI was also not significantly correlated with slow wave sleep (SWS) VmHRV during the consolidation night or wake VmHRV. Finally, no significant correlations were observed between ERI and other REM measures during the consolidation night, including REM%, REM density, REM fragmentation, REM latency, and REM theta.


1. Milad, M.R., et al., *Recall of fear extinction in humans activates the ventromedial prefrontal cortex and hippocampus in concert.* Biol Psychiatry, 2007. **62**(5): p. 446-54.
2. Milad, M.R., et al., *Deficits in conditioned fear extinction in obsessive-compulsive disorder and neurobiological changes in the fear circuit.* JAMA Psychiatry, 2013. **70**(6): p. 608-18; quiz 554.
3. Dawson, M.E., A.M. Schell, and D.L. Filion, *The electrodermal system. In: Cacioppo JT, Tassinary LG, Berntson GG, eds. Handbook of Psychophysiology*. Third ed. 2007, New York, NY: Cambridge University Press.
4. Lonsdorf, T.B., et al., *Don't fear 'fear conditioning': Methodological considerations for the design and analysis of studies on human fear acquisition, extinction, and return of fear.* Neurosci Biobehav Rev, 2017. **77**: p. 247-285.

5. Orr, S.P., et al., *De novo conditioning in trauma-exposed individuals with and without posttraumatic stress disorder.* J Abnorm Psychol, 2000. **109**(2): p. 290-8.
6. Berry, R.B., et al., *The AASM Manual for the Scoring of Sleep and Associated Events: Rules,Terminology and Technical Specifications.* . 2015, Darian, IL: American Academy of Sleep Medicine.
7. Lonsdorf, T.B., C.J. Merz, and M.A. Fullana, *Fear Extinction Retention: Is It What We Think It Is?* Biol Psychiatry, 2019. **85**(12): p. 1074-1082.
